# Experimental challenge of Chinook salmon with *Tenacibaculum maritimum* and *Tenacibaculum dicentrarchi* fulfils Koch’s postulates

**DOI:** 10.1101/2024.03.06.583827

**Authors:** Karthiga Kumanan, Jeremy Carson, Ryan B. J. Hunter, Anne Rolton, Ulla von Ammon, Chaya Bandaranayake, Connie Angelucci, Richard N. Morrison, Seumas P. Walker, Jane E. Symonds, Kate S. Hutson

## Abstract

The bacterial skin disease tenacibaculosis, caused by *Tenacibaculum* species, can compromise numerous species of economically important marine fish, including salmonids. While tenacibaculosis is a known threat to Atlantic salmon (*Salmo salar*) aquaculture, the pathogenesis of *Tenacibaculum maritimum* and *Tenacibaculum dicentrarchi* on Chinook salmon (*Oncorhynchus tshawytscha*) has not yet been investigated. In this study, three molecular O-AGC types of *T. maritimum* (O-AGC Type 3-0, Type 2-1 and Type 3-2) and *T. dicentrarchi* isolated during a disease outbreak of farmed Chinook salmon in Aotearoa New Zealand were assessed for their ability to induce tenacibaculosis in salmon smolts under controlled conditions. Naive Chinook salmon were exposed to *T. maritimum* or *T. dicentrarchi* by immersion. Clinical signs of tenacibaculosis were apparent post-exposure and observed in 100% of all three molecular O-AGC types of *T.-maritimum*-challenged fish, with 100% morbidity in O-AGC Type 2-1 and Type 3-2 and 60% in O-AGC Type 3-0. Chinook salmon exposed to *T. dicentrarchi* showed characteristic clinical signs of disease in 51% of the challenged population, with 28% morbidity. Common gross pathological signs observed for both *Tenacibaculum* species were congruent with observations on farmed fish in the field, including scale loss, erythematous skin lesion, skin ulcers, fin necrosis, mouth erosion and gill ulceration. Exophthalmia was observed only in *T. maritimum*-challenged fish, while skin ulcers appeared grossly more severe with exposed musculature in *T. dicentrarchi*-challenged fish. Pure *T. maritimum* and *T. dicentrarchi* cultures were reisolated from the skin and gills of the challenged fish and their identity was confirmed by species-specific PCR and molecular O-AGC typing. Challenge experiments and associated field surveillance (for *T. maritimum*) did not show the presence of culturable *T. maritimum* cells in the anterior kidney. This provides compelling evidence that tenacibaculosis in farmed Chinook salmon is an external infectious disease, and that *Tenacibaculum* is a marine obligate organism that is unable to survive in fish body fluids and does not cause septicaemia. This has repercussions for approaches to experimental challenges with *Tenacibaculum* species, which must occur by immersion rather than intraperitoneal or intramuscular inoculation, to replicate the natural transmission pathway and to ensure a successful challenge model. This study fulfilled modernised Koch’s postulates for the three molecular O-AGC types of *T. maritimum* and single strain of *T. dicentrarchi* as aetiological agents of tenacibaculosis in Chinook salmon that cause mortalities with considerable external abnormalities.

**Author summary:** Chinook salmon, *Oncorhynchus tshawytscha*, is the most significant species of Pacific salmon for its large size and nutritional content which makes it a premium choice for aquaculture. In Aotearoa|New Zealand, Chinook salmon is the only marine salmon species farmed. For a decade, the industry was impacted by an undiagnosed skin disease resulting in high mortalities. Disease susceptibility in Chinook salmon is scarcely studied and added to the challenge for a timely diagnosis. This novel research provides insight on disease susceptibility of Chinook salmon and confirms *Tenacibaculum* species identified in New Zealand pose a high threat to the aquaculture industry. This research has global implications and contributes valuable insights and approaches to disease management that can be applied in British Columbia and Canada where Chinook salmon populations are in decline.

## 1. Introduction

Salmon are the highest value marine fish species in the global aquaculture sector [1, 2]. Since the 1980s, salmonid aquaculture has been hampered by tenacibaculosis (formerly known as marine flexibacteriosis), an economically significant marine infectious disease [3-7]. Tenacibaculosis is an epidermal infection caused by the biofilm-producing, Gram-negative obligate marine bacteria *Tenacibaculum* spp. on external mucosal surfaces of fish including the skin, fins, gills, palate and teeth [8-12]. *Tenacibaculum* species that are known to cause skin disease of fish include *T. maritimum* [see 5, 8, 13, 14-16], *T. dicentrarchi* [see 16, 17], *T. soleae* [see 18, 19], *T. discolor* [20], *T. piscium* [see 21], *T. bernardetii* [see 22] and *T. finnmarkense* [see 16, 23]. While Atlantic salmon (*Salmo salar*), aquaculture has been impacted by tenacibaculosis since the 1990s [14, 24], farmed Chinook salmon (*Oncorhynchus tshawytscha*) was considered to be resistant to *T. maritimum* in the Pacific Northwest [USA and Canada; 8]. No major tenacibaculosis outbreaks have been reported even though *T. maritimum* was initially detected in Chinook salmon in marine pens in California in the early 1990s [25]. More recently, Bass et al. [26] hypothesised that *T. maritimum* was associated with low survival and poor body condition in wild Chinook and Coho salmon in British Columbia sampled early in their marine life.

*Flexibacter* (most likely *T. maritimum*) was first reported in Aotearoa New Zealand Chinook salmon in 1989 [27]. At the time, a *Flexibacter* species was found in association with skin lesions and significant levels of mortality of juvenile salmon following transfer to marine waters. Since 2012, Chinook salmon farms located in the Marlborough Sounds have consistently reported elevated levels of mortality associated with skin disease. Disease investigations (from 2015 to 2019) of fish exhibiting skin ulcers showed inconsistent detection of *T. maritimum* and/or *Rickettsia*-like organisms by polymerase chain reaction (PCR), indicating a multi-agent disease condition [28-30]. However, in the summer of 2020, a field study, using culture methods, revealed 100% association of *T. maritimum* in clinically compromised fish with skin lesions [31] and no detection of *Rickettsia*-like organisms (NZ RLO1 and NZ RLO2) by PCR. There were also incidences of *T. maritimum* co-infection with *T. dicentrarchi* or *T. soleae* [see 31]. Further analyses revealed that the *T. maritimum* associated with the mortality event represented three molecular O-antigen gene cluster (O-AGC) types (O-AGC Type 3-0, O-AGC Type 2-1 and O-AGC Type 3-2) [32]. Consequently, tenacibaculosis was categorised as a priority disease by the Chinook salmon industry in Aotearoa New Zealand.

The host-pathogen relationship and pathogenesis of tenacibaculosis in Chinook salmon has scarcely been studied. Proving causality of a disease in uncontrolled aquatic environments can be complex due to potential multi-factor influences (i.e. increasing sea surface temperatures and multiple disease agents). In the case of salmon skin disease and mortalities in Aotearoa New Zealand, the scenario is particularly complex given the identification of multiple contributing factors including bacterial pathogens (NZ RLOs, *Tenacibaculum* spp., *Vibrio* spp.) and environmental factors such as warming seawater, harmful algae and biofouling organisms [33-35]. The Henle–Koch postulates [36, 37] to prove causality of disease have been successfully applied to several aquatic diseases [see 38 for review]. Some examples include shrimp white faeces syndrome [39], francisellosis in Atlantic cod (*Gadus morhua*) [see 40], amoebic gill disease in Atlantic salmon [41], bacterial and fungal diseases in marine seaweed [42, 43], streptococcosis in whiteleg shrimp (*Litopenaeus vannamei*) [see 44], and heart- and skeletal muscle inflammation in Atlantic salmon [45]. While Koch’s postulates have been fulfilled for *T. maritimum* and *T. dicentrarchi* in Atlantic salmon [5, 8, 16], the susceptibility of Chinook salmon to tenacibaculosis has not been demonstrated experimentally.

The aim of this study was to establish whether *T. dicentrarchi* and all three O-AGC types of *T. maritimum*, isolated from Aotearoa New Zealand Chinook salmon, can cause skin disease in Chinook salmon in fulfilment of Koch’s postulates. A second aim was to investigate whether *Tenacibaculum* species can proliferate within internal organs of fish and cause septicaemia. Several published studies have suggested that *T. maritimum* can cause systemic infections [8, 16, 24, 46]. However, we hypothesised that *T. maritimum*, being an obligate marine organism, requires sodium levels to be approximately equimolar to seawater for its viability. This characteristics would inhibit replication of the pathogen within the body of fish where osmoregulation reduces plasma osmolarity within a range of 290-340 mOsmol/L [47, 48], which approximates to a sodium chloride concentration of 0.9% w/v. We obtained field data from fish exhibiting clinical tenacibaculosis on farm and laboratory derived data from the challenge trials to assess the presence of viable *T. maritimum* in the kidneys of affected fish.

## 2. Materials and methods

### 2.1. Animal ethics statement

All procedures performed involving live salmon were carried out in accordance with New Zealand government regulations and associated ethical standards as administered by the Nelson Marlborough Institute of Technology Limited Animal Ethics Committee approval (Ref. AEC2018 CAW011 and AEC-2022-CAW-02).

### 2.2. Source of fish and husbandry

Juvenile Chinook salmon (*Oncorhynchus tshawytscha*) were sourced from a freshwater hatchery at Tentburn, New Zealand. Fish had been vaccinated for *Yersinia ruckeri* only. Salmon were transported in fresh water in a custom-built transport vehicle with oxygen control, off-gassing and real-time monitoring. The salmon were housed in 1,000 L tanks at *Te Wero Aro-anamata*, Cawthron’s aquatic physical biocontainment (PC2) facility. Each tank was equipped with its own recirculating aquaculture system (RAS), which consisted of temperature control ±0.3°C through a supervisory control and data acquisition system, real time alarmed oxygen monitoring, particle filtration with bubble bead filters, inline ultraviolet (UV) disinfection at 56 mj/cm^2^ dose and 50 L/min water flow, foam fractionator for fine particle removal and a biofilter. Natural seawater used in this study was sourced from the Cawthron Aquaculture Park, Glenduan, Nelson, New Zealand, and was UV treated and filtered to 5 μm prior to use. On arrival, the fish were transferred into water at 17°C and 15 ppt salinity and were then gradually acclimated to 35 ppt water salinity over 16 days. The fish were fed to satiation three times per day. Fish were not fed for 2 days prior to manipulation and for the duration of the experiment.

### 2.3. Bacterial strain and propagation

*Tenacibaculum maritimum* strains in Aotearoa New Zealand belong to three O-AGC types [Cawthron Culture Collection of Microorganism (CCCM20/006, CCCM20/102 and CCCM20/133) see Table 1; 32]; *Tenacibaculum dicentrarchi* (CCCM21/136) used in this study were isolated from moribund Chinook salmon skin ulcers during a disease outbreak in the Marlborough Sounds, Aotearoa New Zealand. The isolates were confirmed to be *T. maritimum* and *T. dicentrarchi* by species specific PCR [31, 49, 50]. The isolates were stored at -80 °C on cryobeads (Microbeads^TM^) and revived on Marine Shieh’s Medium agar [MSM, 50] and ZoBell’s Marine Agar 2216E (MA) and then incubated at 22°C for 48 h. Growth of *T. maritimum* on both types of media was assessed and purity checked by Gram-stain. Pure colonies were inoculated into 50 mL primary MSM and MB, respectively, in 150 mL baffled conical flasks and incubated at 22°C for 48 h at 180 revolutions per minute (RPM) on a Ratek orbital shaker. Primary broths were used to inoculate 1 L MSM and MA broth for the challenge trial. An approximate cell density was assessed by microscopy using a Helber bacteria counting chamber. To assess cell viability by plate count, 100 μL of tenfold dilution of broth culture was inoculated onto MSM using a spread plate method and incubated at 22°C for 48 h. Colony-forming units (CFU) were enumerated to determine viable cell concentration in the immersion challenge.

**Table 1.**
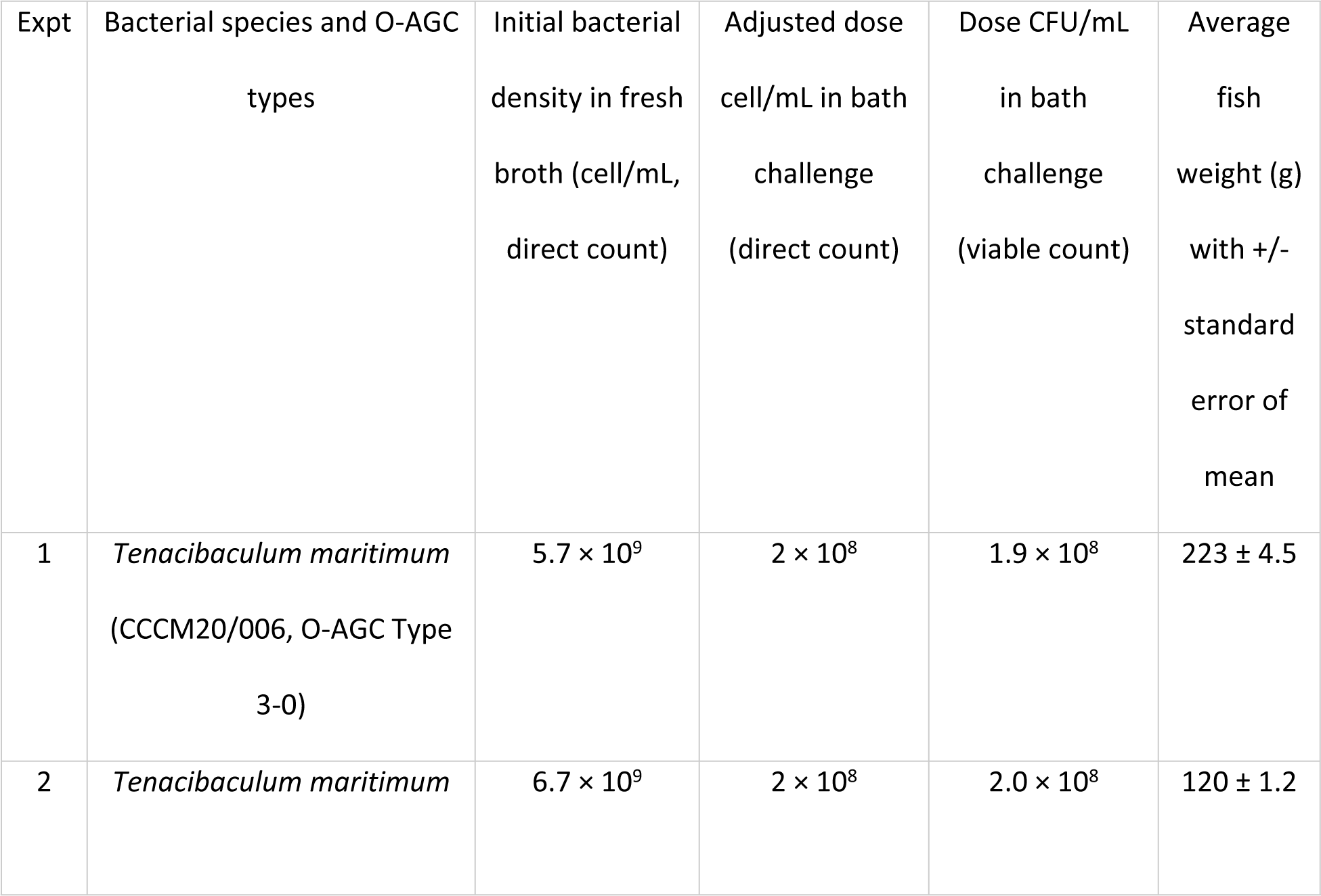
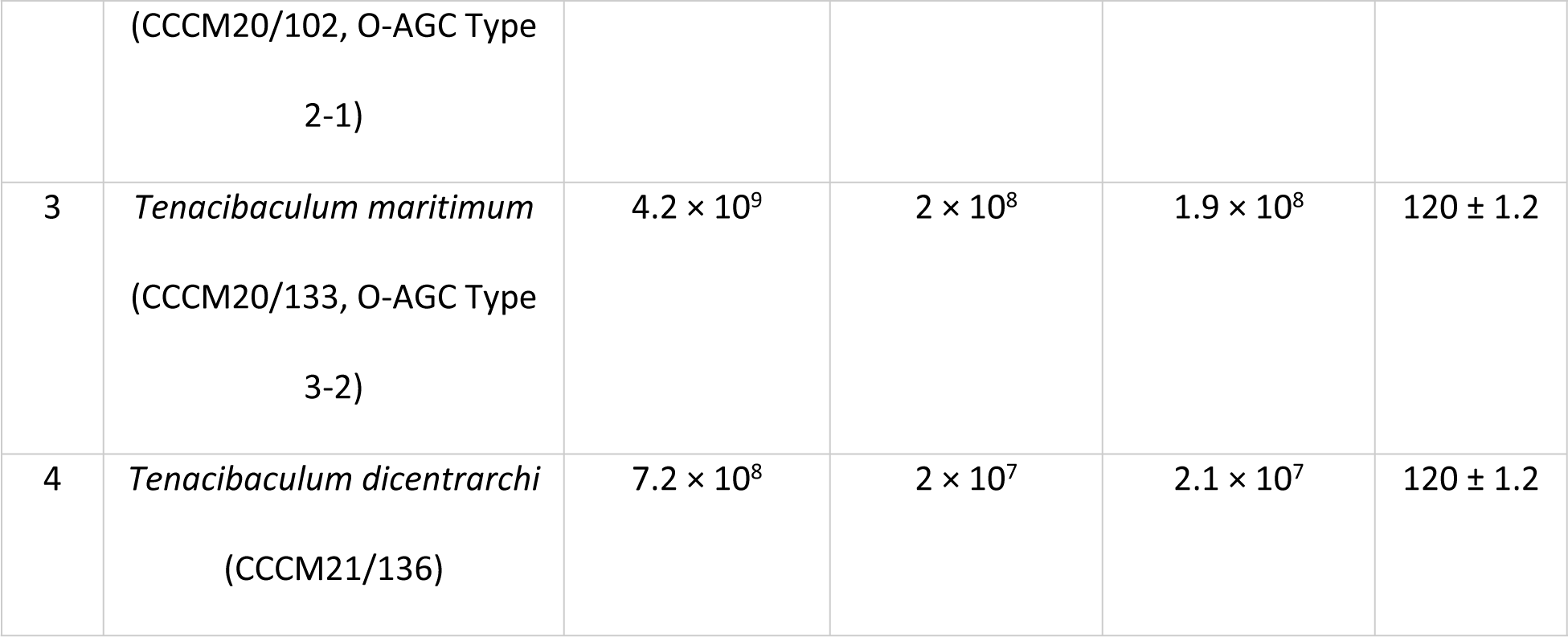
*Tenacibaculum* species and strains used in the Chinook salmon experimental challenge trials indicating bacterial concentration used in each challenge. Initial bacterial density in fresh broth = bacterial density obtained post incubation; adjusted dose cell/mL in bath challenge = bacterial density used in the challenge bath.

### 2.4. Pre-trial health check of naïve fish and tank water bacteriology

The mean weight of experimental fish used in the *T. maritimum* and *T. dicentrarchi* infection challenges are as shown in Table 1. Five fish per tank were humanely euthanised prior to sampling by percussive stunning. These fish were sampled for bacteriology for potential pathogenic bacterial species such as *T. maritimum* and *T. dicentrarchi* and molecular analysis to screen for NZ-RLO1, NZ-RLO2 and *Y. ruckeri*. Euthanized fish were placed on a sterile surface. Skin mucus and a gill swab were collected for each fish using a sterile swab and directly streaked onto Marine Shieh’s Selective Medium (MSSM) [50], thiosulfate–citrate–bile salts–sucrose agar (TCBS), MA and Columbia sheep’s blood agar (5% blood and 0.5% NaCl) (BA). The fish were aseptically dissected to expose the internal organs. Using sterile forceps and a scalpel blade, the digestive system was removed, without puncturing the digestive content or the heart, to expose the anterior kidney. The epithelial membrane covering the anterior kidney was removed using a sterile scalpel blade and a sterile swab was inserted directly into the anterior kidney, avoiding contact with any other internal organs. The swab was directly inoculated onto all four media as described. All inoculated media were incubated at 22°C for 72 h. Different sets of sterile tools were used for each fish sampled to avoid cross-contamination of the anterior kidney with external bacterial flora. Approximately 3 mm^3^ of skin, gill and anterior kidney tissue were aseptically dissected and stored in DNA/RNA Shield (Zymo Research) at 4°C for molecular analysis. A tetraplex assay was employed to detect and quantify the four primary pathogens of Aotearoa New Zealand Chinook salmon following von Ammon et al. [49]. Fish holding RAS tank water was tested for the presence/absence of *Tenacibaculum* species by plating 100 μL of water onto MSSM and incubated at 22 °C for 72 h.

### 2.5. Experimental infection of *T. maritimum* and *T. dicentrarchi* by immersion

*Tenacibaculum maritimum* isolates of the three O-AGC types [O-AGC Type 3; Type 2-1 and Type 3-2; 32] and *T. dicentrarchi* grown in MSM broth were used for separate immersion challenges at a fish density of 60 kg/m^3^ in the immersion bath. For each trial, two experimental groups (control and treatment) were tested in triplicate (Figure 1). The experimental challenge was performed at a water temperature of 17 °C. Control (n = 30 fish) and treatment groups (n =30 fish) were transferred by net to individual 100 L polypropylene fish bins containing UV-treated natural seawater at 35 ppt salinity. Oxygen (100–110% saturation) was supplied to each challenge bath during the pathogen challenge. Control and treatment challenge baths were separated by at least 20 m to avoid cross contamination. The control and treatment groups were challenged simultaneously. The bacterial concentration in the bath was standardised to 2 x 10^8^ cells/mL for all three O-AGC types of *T. maritimum* and 2 x 10^7^ cells/mL of *T. dicentrarchi* as described by Nowlan et al. [16]. *Tenacibaculum dicentrarchi* bacterial density in the challenge bath was tested at 2 x 10^7^ cells/ml due to the lower bacterial yield in broth culture than that achieved for *T. maritimum*. The same volume of sterile MSM broth was added to UV treated seawater in the control bath for the challenges. A lid was placed on top of the bin to minimise splash fall-out and fish were held immersed for 60 min. Fish from the treatment and control groups were redistributed to individual triplicate RAS tanks stocked with 10 fish/tank for all *T. maritimum* challenges and 13 fish/tank for the *T. dicentrarchi* challenge (Figure 1). Fish were monitored every 3 h for clinical and behavioural signs of disease such as scale loss, skin lesions, skin ulcers, fin erosions, epidermal haemorrhage, exophthalmia and mouth erosion.

**Figure 1.**
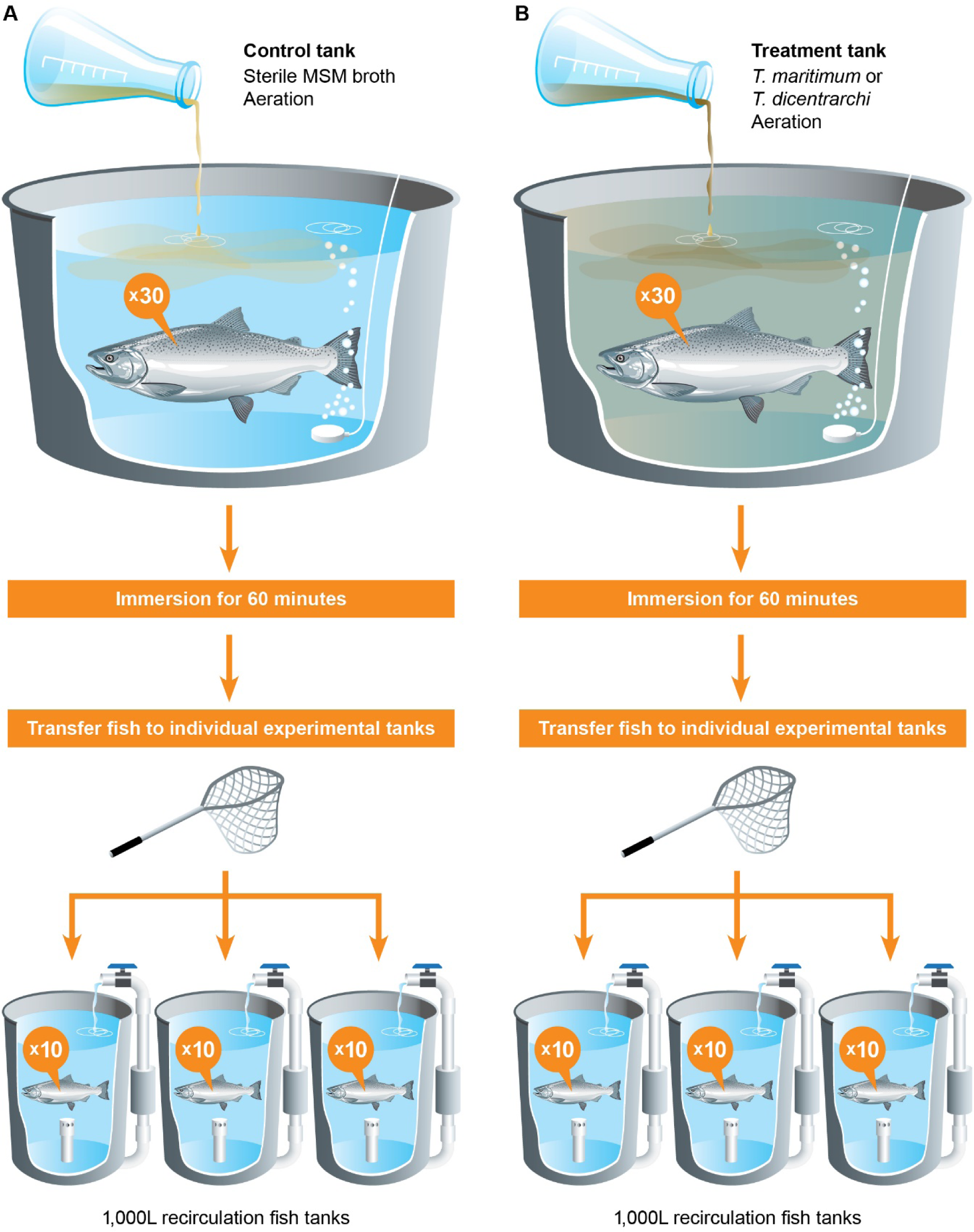
Schematic representation of the *in vivo* immersion challenge of Chinook salmon with three *Tenacibaculum maritimum* strains (O-AGC Type 3-0, Type 2-1 and Type 3-2) and *T. dicentrarchi*.

### 2.6. Post-mortem and end of trial examination

Taking fish welfare into consideration as we observed disease progression, the *T. maritimum* O-AGC Type 3) and *T. dicentrarchi* challenge experiments were terminated at 8 and 11 days post-infection (DPI), respectively. *Tenacibaculum maritimum* trials of strain O-AGC Type 2-1 and Type 3-2 were ended at 3 DPI. Throughout the experiment, fish found moribund or dead were immediately removed for necropsy. The survivors from the treatment and control groups at the end of experiment were euthanized by iki-jime [51]. Fish from the control group were euthanized and sampled first, followed by the treatment group, to prevent contamination.

#### 2.6.1. Bacteriological analysis

Fish were placed sample side up (side presenting clinical disease), and sampled only on one side, to avoid contamination from the supporting surface. Bacterial swab samples were collected from external lesions and the anterior kidney. Bacterial swab samples collected from the skin, mouth, gill or tail lesions were inoculated onto MSSM to reisolate the organism used in the challenge. To rule out the presence of other pathogens that cause skin disease, samples were also inoculated on TCBS, BA and MA, and incubated as described in Section 2.4 for a minimum of 4 days. After incubation, bacterial colony morphology consistent with *T. maritimum* and *T. dicentrarchi* was assessed on MSSM and MA [see Figure 2 in 31]. Species identification and the O-AGC types were confirmed for the bacterial colonies using species specific PCR and mPCR [32, 49, 50, 52]. In addition, to investigate whether *T. maritimum* or *T. dicentrarchi* causes systemic infection, a swab from the anterior kidney was plated onto all four media types as described in Section 2.3. Viable *Tenacibaculum* counts in RAS tank water were quantified daily for all trials from 1 DPI to the end of a trial for each *Tenacibaculum* species /strain. One hundred microlitres of tenfold diluted RAS tank water was plated onto MSSM in triplicate and incubated at 22°C for 72 h.

**Figure 2.**
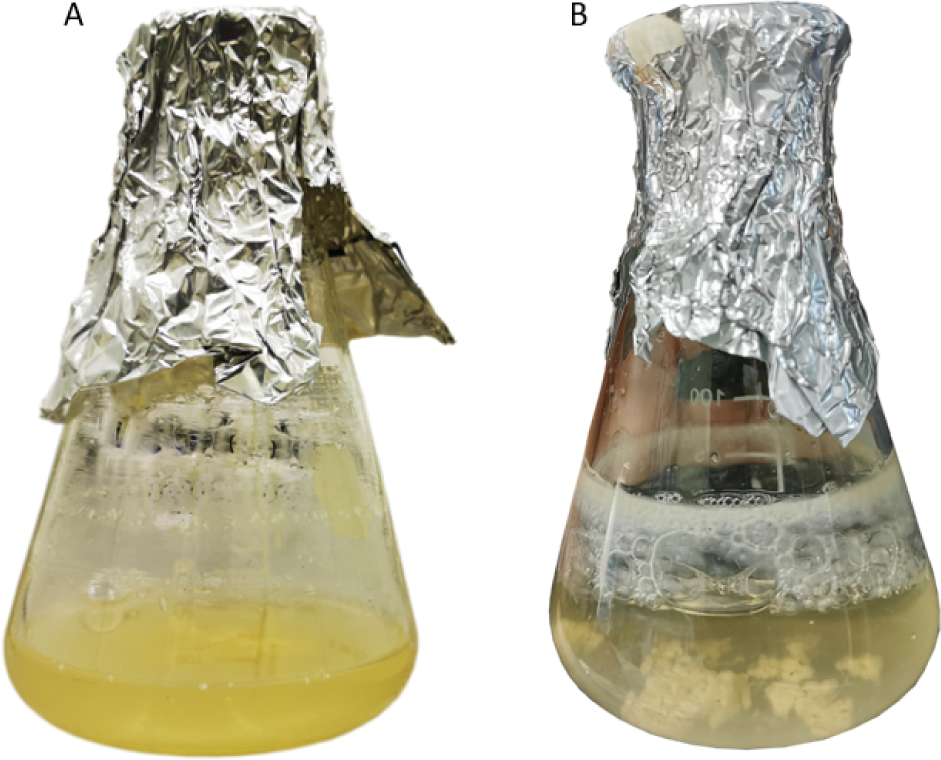
*Tenacibaculum maritimum* culture (O-AGC Type 3-0) incubated for 48 h at 22 °C in MSM broth (A) and marine broth (ZoBell 2216E; B). Culture in MSM broth yielded higher growth with no aggregation compared to the culture grown in marine broth.

#### 2.6.2. Histopathology

Histopathological analysis was carried out on fish from *T. maritimum* (O-AGC Type 3-0, n =7) and *T. dicentrarchi* (n = 3) and control (n = 3) trials. Approximately 5 mm^3^ of skin tissue (periphery of the lesion), gill and anterior kidney were excised and preserved in 1:10 parts of 10% neutral buffered formalin. Tissue samples were sent to Gribbles Veterinary Pathology, Christchurch, New Zealand for processing, paraffin embedding and haematoxylin and eosin (H & E) staining. The authors performed histopathological examination to evaluate the presence or absence of pathology.

#### 2.6.3. Droplet Digital PCR analysis of infected fish

Tissue samples of fish from *T. maritimum* O-AGC Type 3-0 (control n = 30 and challenge n = 29, one fish was excluded due to bleach contamination) and from *T. dicentrarchi* (control n = 15, challenge n = 15) challenge trials were analysed to investigate presence of DNA from pathogens in various organs of infected fish. Approximately 30 mg of individual fish tissue (skin, gill and anterior kidney) from diseased fish, was used for genomic DNA (gDNA) extraction using Quick-DNA™ Miniprep Plus Kit (Zymo Research) following the manufacturer’s instructions. Tissues from control fish were processed prior to samples from the treatment group and each tissue type was processed on separate days to avoid potential cross-contamination during DNA extraction. One hundred nanograms of gDNA from each tissue type or isolate were used to perform ddPCR assay using a QX200 Droplet Digital PCR System^TM^ (Bio-Rad, Hercules, USA), as described by von Ammon et al. [49]. *Tenacibaculum dicentrarchi*-specific primers and PCR conditions as described in Wilson et al. [50] were used in droplet digital PCR to quantify *T. dicentrarchi* DNA in fish tissue infected with *T. dicentrarchi*.

#### 2.6.4. Systemic vs external infection investigation

The ability for *T. maritimum* to survive and proliferate within the internal organs of Chinook salmon was investigated on diseased fish sampled from commercial salmon farm sites in the Marlborough Sounds, South Island (Otanerau, *n* = 28, Waihinau, *n* = 8 and Kopāua *n* = 5), and from salmon used in the challenge experiment. All fish exhibited various clinical signs of tenacibaculosis including scale loss, skin ulcers, erythematous skin lesion, and fin and mouth erosion. We used a bacteriological method to detect the presence or absence of viable *T. maritimum* and compared it against positive ddPCR detection, which amplifies nucleic acid from both viable and non-viable cells. Fish from the farm were transported to Cawthron’s diagnostic laboratory in separate plastic bags on ice, and a necropsy was carried out within 12 h. A swab sample from the skin and anterior kidney was inoculated onto MSSM to compare the presence of viable *T. maritimum* between external and internal tissue. Fish necropsies as described in Section 2.4, were performed with utmost care to avoid contaminating the anterior kidney with external bacterial flora present on fish skin. Molecular analysis of *T. maritimum* was conducted on anterior kidney as described in Section 2.6.3.

### 2.7. Statistical analysis

#### 2.7.1. Kaplan-Meier survival curve

R-Studio version 4.2.1 [53] was used for all statistical analyses. Survival of Chinook salmon in each experimental group was plotted using the Kaplan-Meier survival curve with the R package ‘survival’ and applying the daily morbidity data.

#### 2.7.2. Statistical analysis of bacterial load and correlation with health status

Differences in ddPCR detection of *T. maritimum* O-AGC Type 3 and *T. dicentrarchi* in copies/μl and square root-transformed for the factor ‘tissue’ (skin, gill, and anterior kidney) and the second factor “health status” (mortality, moribund and survivors) were analysed within the R Project statistical software [53]. The non-parametric two-factorial Scheirer– Ray– Hare tests were undertaken, followed by post hoc pairwise Wilcoxon comparisons, considering results as significant if *p* ≤ 0.05. Survival in the control groups was 100% (unaffected); therefore these were excluded from statistical analysis. In addition, linear regression curves were plotted to compare bacterial loads (copies/µl) between the anterior kidney and the combined external tissues (skin and gill) for *T. maritimum* and *T. dicentrarchi*, respectively. A test for correlation (using Pearson’s correlation coefficient) was used to test for significance.

## 3. Results

### 3.1. Bacterial strain and propagation and pre-trial assessment

Three *Tenacibaculum maritimum* strains (O-AGC Type 3-0, Type 2-1 and Type 3-2) revived from micro beads performed consistently better on MSM agar and in broth compared to Marine Agar and broth. MSM resulted in higher cell densities, less aggregation, and less biofilm production (Figure 2).

*Tenacibaculum dicentrarchi* did not form aggregates in either medium, and the cell density in MSM broth was higher compared to MB. *Tenacibaculum maritimum* and *T. dicentrarchi* cultured in MSM broth with vigorous shaking at 180 rpm resulted in a yield of 10^9^ cells/mL and 10^8^ cells/mL, respectively, by direct count using the Helber counting chamber (Table 1). Bacterial concentration by direct count for strains of *T. maritimum* and *T. dicentrarchi* were then adjusted to 2 x 10^8^ cell/mL and 2 x 10^7^ cell/mL, respectively, for the bath challenge. However, the viability assessment performed on the final challenge bath showed ± 10% variation in the viable bacterial concentration in comparison to direct cell count (Table 1).

The pre-trial fish health assessment confirmed that the fish were in optimal health at the time of challenge and free from all the tested pathogens. Bacterial analysis conducted on tank water confirmed that there were no pre-existing *Tenacibaculum* species in the RAS.

### 3.2. Clinical signs and gross pathology

#### 3.2.1.Tenacibaculum maritimum challenge

Fish were transferred to the individual RAS tanks following 1 h of immersion challenge. On 1 DPI, fish challenged with O-AGC Type 3-0, 2-1 and 3-2 in independent trials were observed to have a reduced swimming rate and displayed white lesions on their abdomen. On close observation, it was confirmed that the patches were scales protruding from the epidermis. At 2 DPI in the *T. maritimum* O-AGC Type 3-0 challenge, erratic swimming and loss of equilibrium was observed in moribund fish and the first mortality occurred. Increased opercular activity was common and the cumulative survival of Chinook salmon in the O-AGC Type 3-0 treatment group at 8 DPI was 40 ± 20% (Figure 3A). On the other hand, smaller fish challenged with strains of O-AGC Type 2-1 and 3-2 reached 100% mortality within 60 h post-challenge (Figure 3B & 3C).

**Figure 3.**
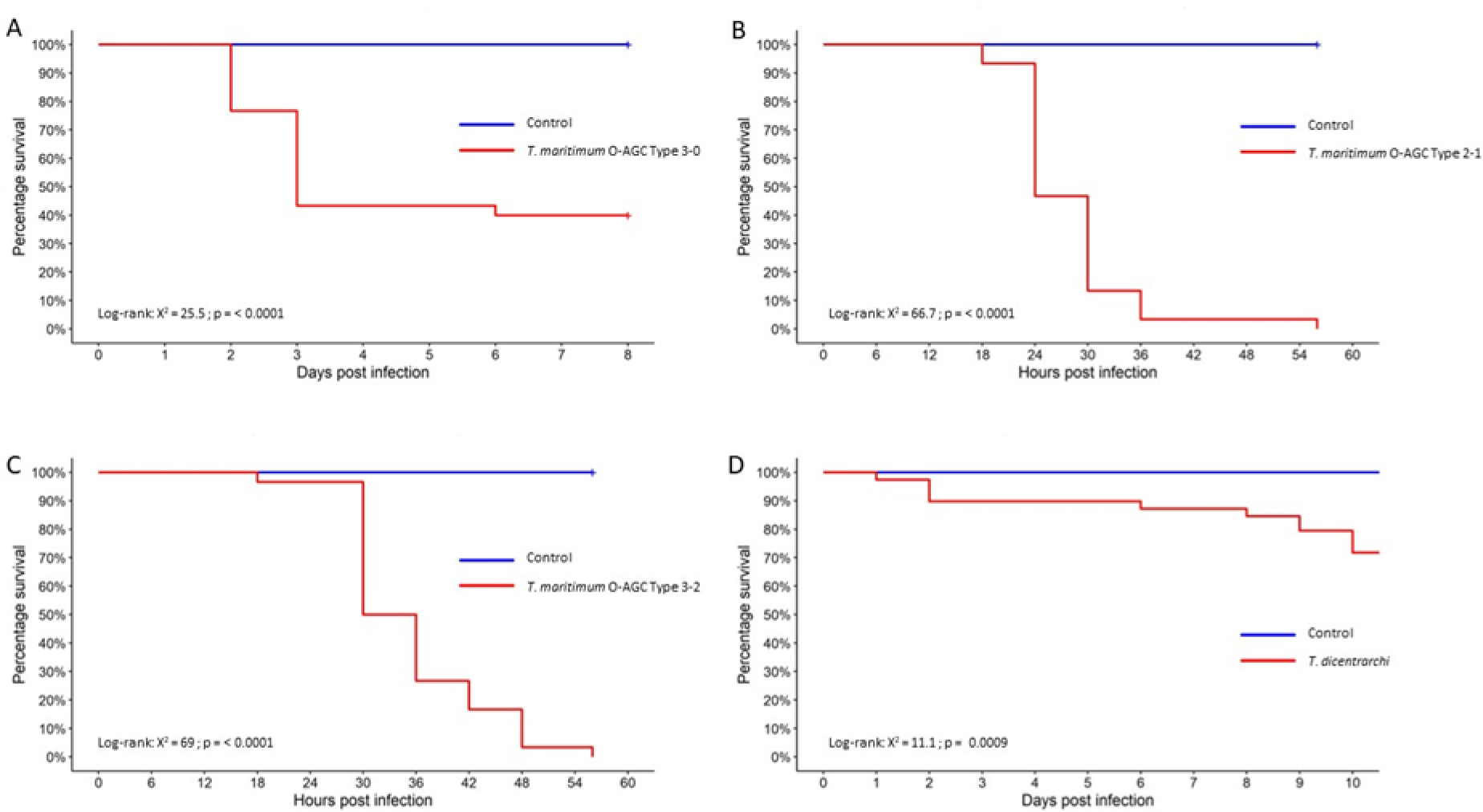
Kaplan-Meier survival analysis of naïve Chinook salmon *Oncorhynchus tshawytscha* experimentally challenged with three molecular O-AGC type strains (A. CCCM20/006, O-AGC Type 3-0; B. CCCM20/102, O-AGC Type 2-1; and C. CCCM20/133, O-AGC Type 3-2) and D. *Tenacibaculum dicentrarchi* (CCCM21/136).

Signs of gross pathology consistent with tenacibaculosis were apparent in all fish exposed to all three *T. maritimum* O-AGC types including mortalities, moribund and surviving fish. Affected fish exhibited varying degrees of pathology ranging from scale loss, skin ulcers, erythematous skin lesion, pectoral and tail fin necrosis, mouth erosion, gill erosion, ventral haemorrhaging, and exophthalmia (Figure 4). No consistent pathologies were observed in the internal organs of experimental fish except for one surviving fish from the *T. maritimum* O-AGC Type 3-0 treatment group, which exhibited severe necrosis of the kidney epithelial membrane (Figure 4I; note, no viable *T. maritimum* was recovered from this fish). No gross pathological signs or mortalities were observed in fish from the control groups.

**Figure 4.**
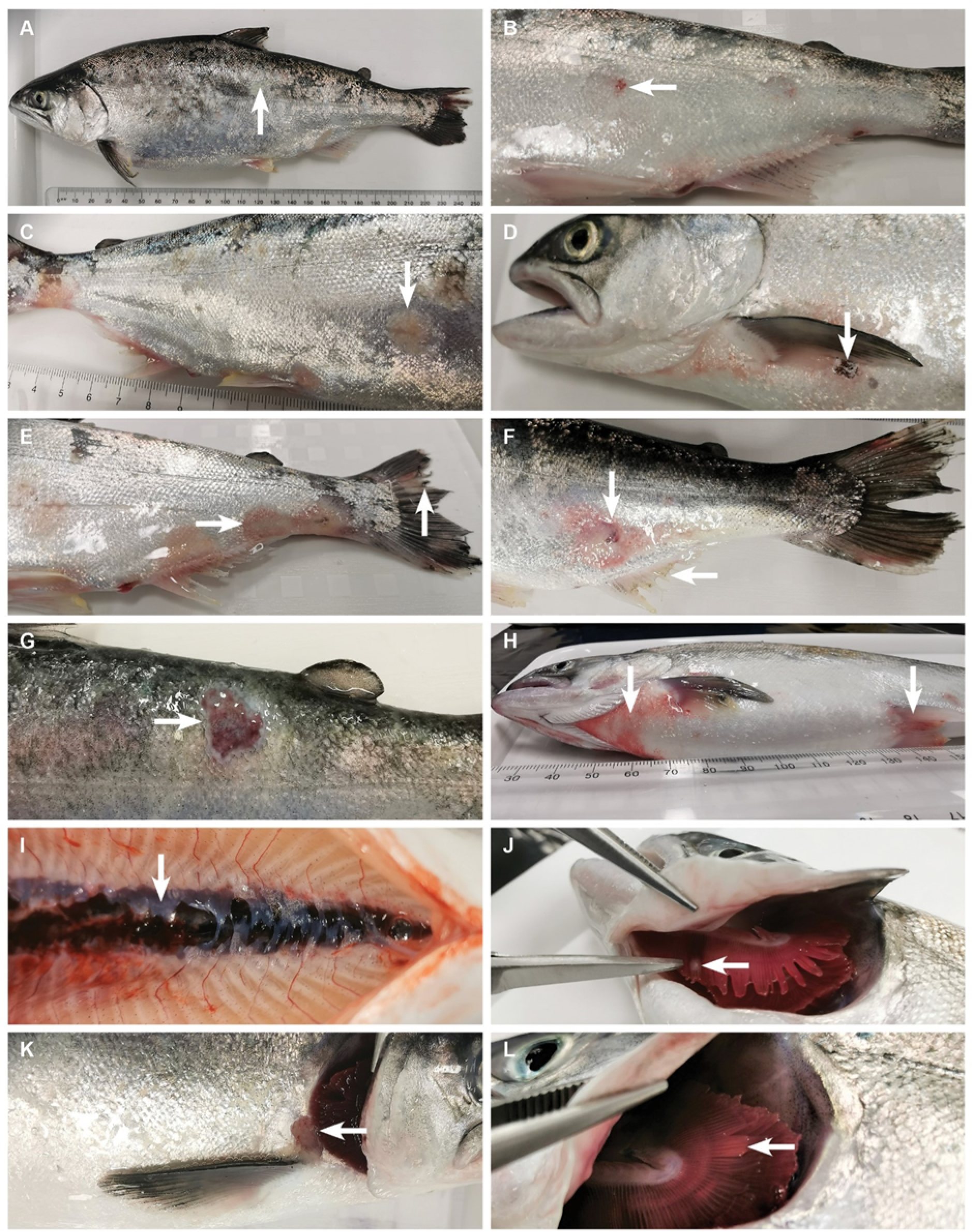
Gross pathology of Chinook salmon *Oncorhynchus tshawytscha* experimentally challenged with *Tenacibaculum maritimum* via immersion. (A) Scale loss. (B) Erythematous skin lesion. (C) Skin lesion with bacterial mat. (D) Skin ulcer under pectoral fin. (E) Tail necrosis and ulcer development towards caudal peduncle. (F) Fin necrosis and abdominal ulcer development. (G) Skin ulcer. (H) Pelvic fin and ventral haemorrhaging. (I) Necrotising anterior kidney epithelial membrane. (J) Yellow plaque on gill filament. (K) Eroded cleithrum bone. (L) Gill erosion.

#### 3.2.2. *Tenacibaculum dicentrarchi* challenge

*Tenacibaculum dicentrarchi* challenged fish started showing increased opercular activity and developed pale white spots on their skin at 3 h post-infection. At 1 DPI, the first mortality was recorded, and individuals developed skin lesions. Common areas of ulceration were under the pectoral fin and the caudal peduncle. Fish exhibited deeper and larger epidermal ulcers, with the musculature exposed, compared with those of the *T. maritimum* challenge which showed more extensive scale loss and less ulceration. A range of pathologies were observed including deep circumscribed ulcers on the abdomen, erythematous skin lesion, mouth rot, spreading skin ulcers with bacterial mats and ulcerated caudal peduncles (Figure 5). Cumulative survival at 11 DPI was 72 ± 9.2% (Figure 3D) with 51 ± 11% of the challenged population exhibiting skin abnormalities as shown in Figure 5. No signs of disease and no mortalities were observed in the control population.

**Figure 5.**
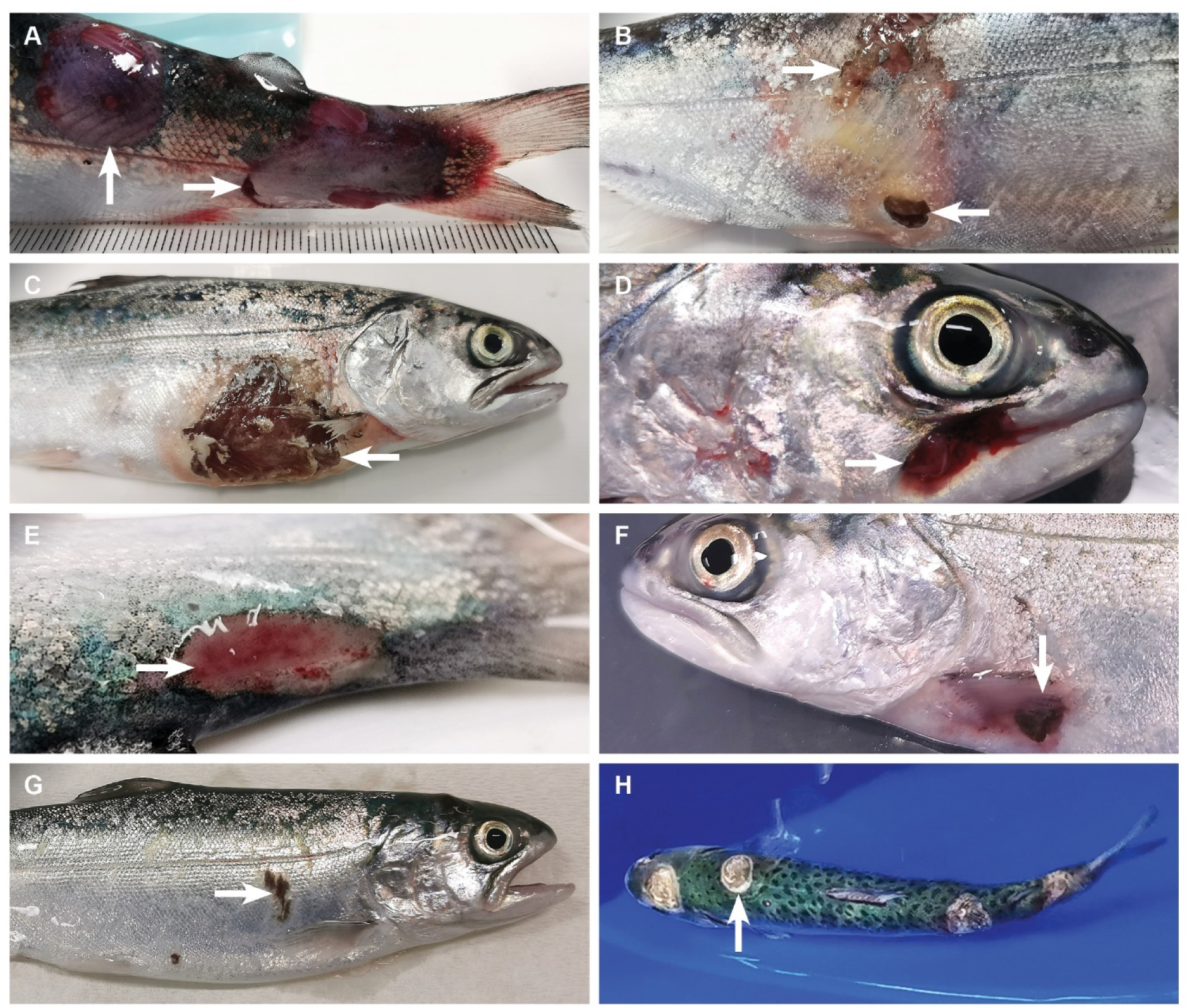
Gross pathology of Chinook salmon (*Oncorhynchus tshawytscha*) experimentally challenged with *Tenacibaculum dicentrarchi* via immersion. (A and E) Epidermal ulceration on the caudal peduncle at 2 DPI. (B) Ulcers developing above the anal fin and expanding to the lateral line with yellow bacterial mat formation. (C and F) Moribund fish exhibiting severe inflammation and ulceration under the pectoral fin. (D) Mouth rot. (G) Skin lesion with intact dermis. (H) Moribund fish prior to removal from tank with multiple circumscribed ulcer patches.

### 3.3. Pathogen identification and fulfilment of Koch’s postulates

#### 3.3.1.Bacteriology

Viable *Tenacibaculum* counts performed on the RAS tank water showed a daily reduction of bacterial load in the experimental tanks despite the disease progression on fish (Figure 6A). All O-AGC types of *T. maritimum* and *T. dicentrarchi* were reisolated from affected fish in the respective challenges. Bacteria were recovered in culture from overtly infected sites including the mouth, skin, and gills. Recovered *T. maritimum* isolates were confirmed to be the respective O-AGC type by mPCR (results not shown). *Tenacibaculum maritimum* and *T. dicentrarchi* were not isolated from the anterior kidney in the challenged population (Figure 6B and 6C) nor from any tissue of the control population.

**Figure 6.**
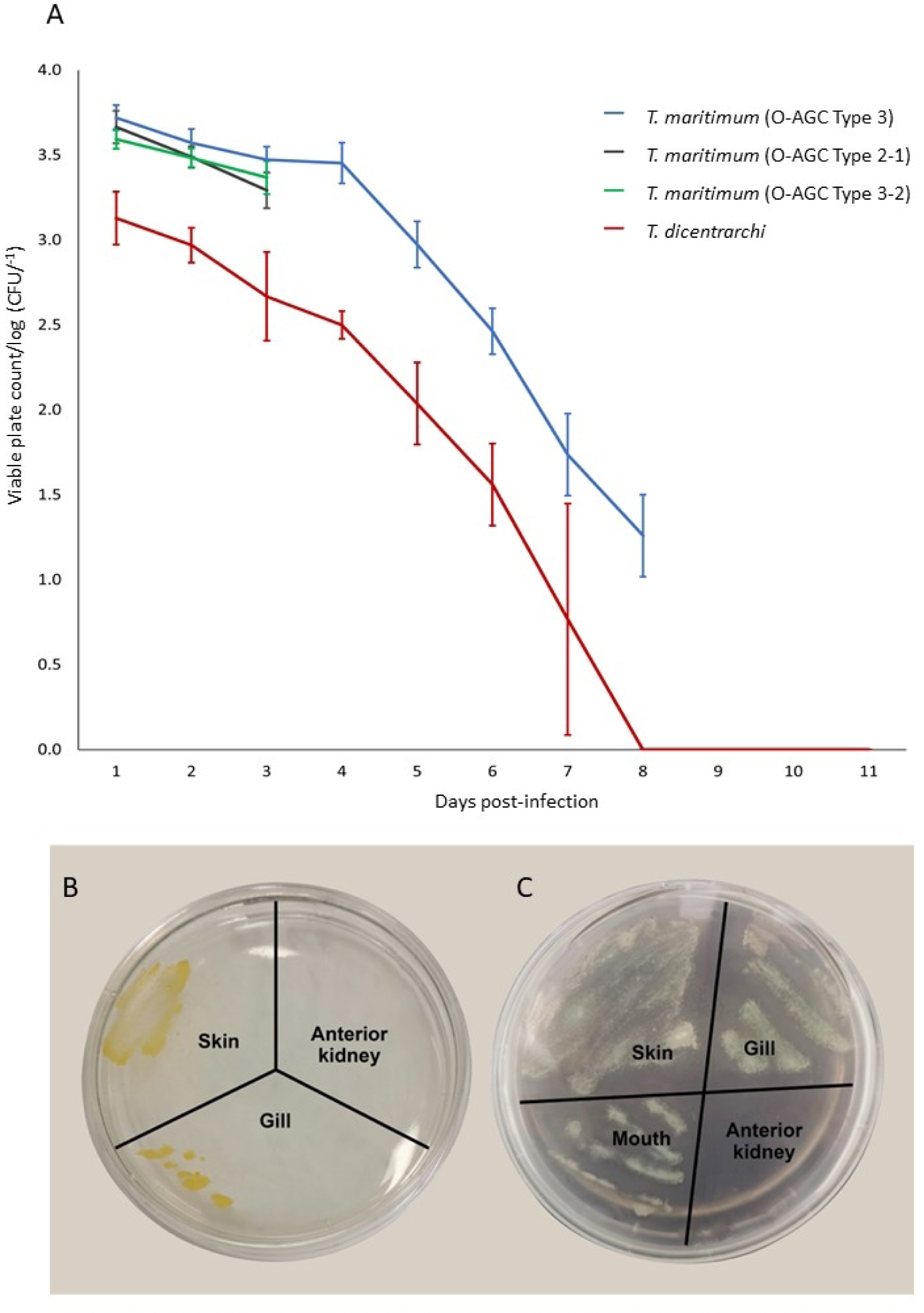
(A) The daily average reduction of viable *Tenacibaculum* cells in RAS tanks during the full experimental duration for each strain. *Tenacibaculum maritimum* strain O-AGC Type 2-1 and Type 3-2 experiments were ended on day 3. (B) *Tenacibaculum dicentrarchi* and (C) *T. maritimum* from infected fish skin, gills, and mouth. The media showed no growth for the sample inoculated from the anterior kidney of the same individual infected fish.

#### 3.3.2. Droplet digital PCR analysis of *T. maritimum* and *T. dicentrarchi* detection in various tissues from infected fish

Samples from control fish in all experimental challenge trials tested negative for *T. maritimum* and *T. dicentrarchi* by ddPCR assay. In fish challenged with *T. maritimum* (O-AGC Type 3-0) and *T. dicentrarchi*, DNA from the pathogens was detected in the skin, gills, and anterior kidneys (Supplementary Table 1). For fish challenged with *T. maritimum*, the highest median gene copy numbers (= 34 copies/µl) were skin samples from moribund and dead fish. Highest outliers of ≥ 75 copies/µl were found in gill samples from moribund and dead fish. For fish challenged with *T. dicentrarchi*, copy numbers were generally lower for all tissues with the highest outliers in gill samples from moribund fish (64 copies/µl) and skin samples from moribund fish (> 60 copies/µl).

Significant differences in *T. maritimum* detection were observed between the sub-categories of the considered health status factors (survivors, moribund and mortality) and tissue types (Scheirer-Ray-Hare, *p* < 0.001; Figure 7). For the *T. dicentrarchi* challenge, the combined factors of fish health status and tissue types were significant (*p* < 0.03). Bonferroni corrected pairwise Wilcoxon rank sum tests showed that gill and skin tissue had significantly higher *T. maritimum* loads compared with anterior kidney (*p* < 0.001), and for mortality and moribund fish compared with survivors (*p* < 0.01). For the *T. dicentrarchi* challenge, the pathogen load as indicated by ddPCR analyses was significantly higher in mortalities compared to survivors (*p* = 0.016). However no significant differences were observed among tissue types (*p* > 0.05).

**Figure 7.**
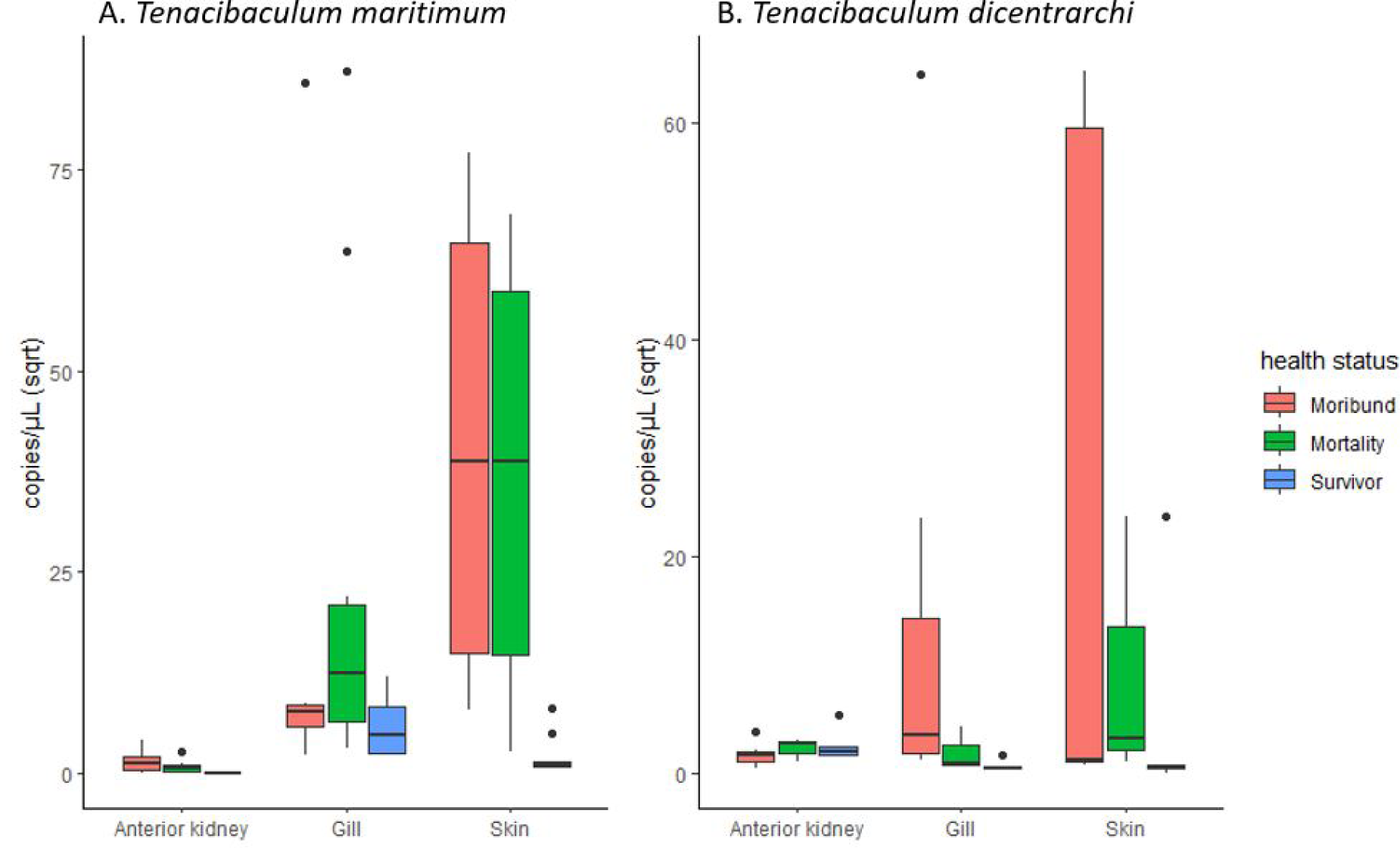
Box plot showing the abundance of (A) *Tenacibaculum maritimum* O-AGC Type 3 and (B) *Tenacibaculum dicentrarchi* (copies/μL (sqrt)) in challenged fish from three categories (moribund, mortality and survivors) and for three tissues (anterior kidney, gills, and skin). (A) Significant difference between anterior kidney and either gill or skin (Bonferroni corrected pairwise Wilcoxon rank sum tests; p < 0.001). Significant difference between survivors and either moribund or mortality fish (stats test; p < 0.01). (B) Significant difference between survivors and mortality (stats test; p = 0.016). Black dots = outliers; Sqrt = Square root transformed.

Bacterial detection between external tissue (skin and gill) versus anterior kidney by ddPCR showed significant positive correlations for *T. maritimum* (copies/mL (sqrt), *p*-value = 0.0066). In contrast, for *T. dicentrarchi* there was no significant correlation between the pathogen load of the internal and external organs (Figure 8).

**Figure 8.**
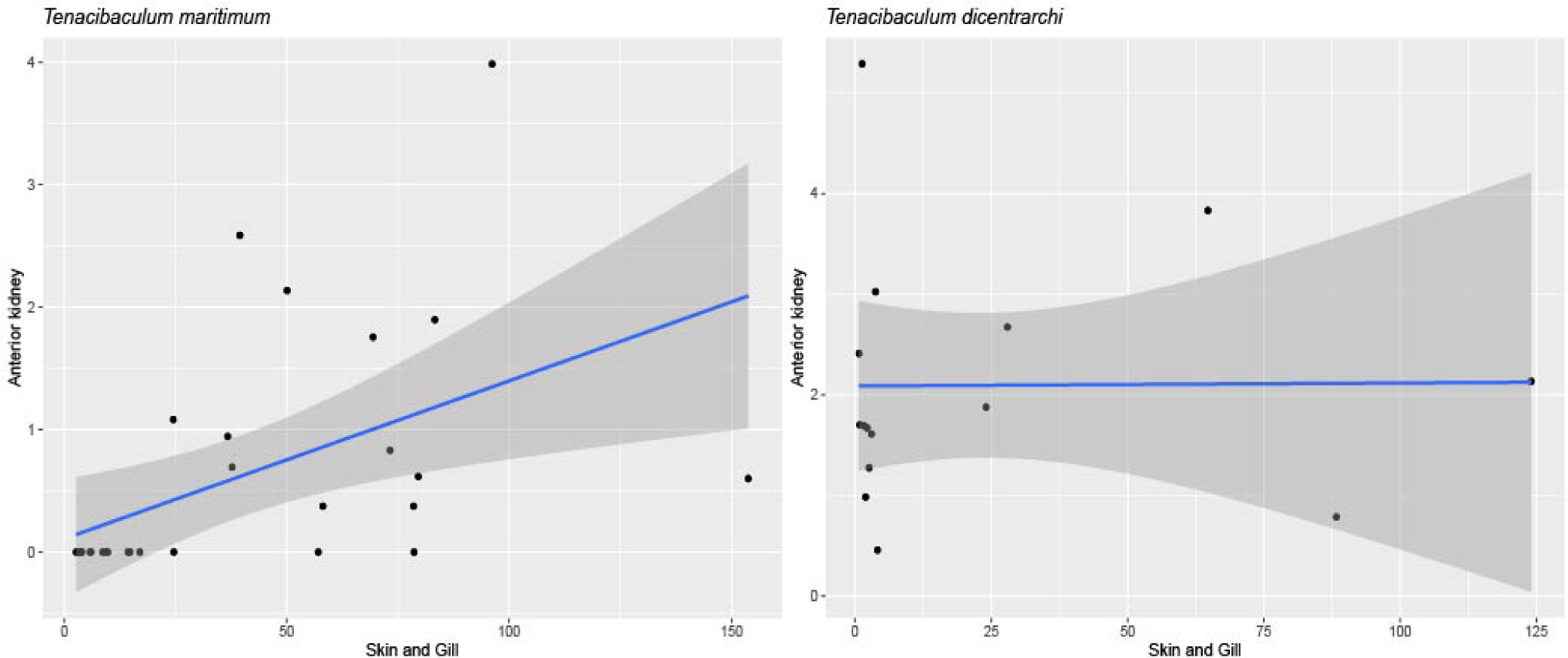
Linear regression curves displaying the correlation of square root transformed pathogen loads (copies/mL) between anterior kidney and external tissue (skin and gill) for *Tenacibaculum maritimum* and *T. dicentrarchi*, respectively.

#### 3.3.3. Histopathology

Histopathological examination of representative samples of infected and uninfected Chinook salmon skin showed distinct differences in the epidermal integrity. The scales and overlying epidermis were absent or were severely damaged in salmon exposed to both *Tenacibaculum* spp. (Figure 9B-F). Fish infected with *T. maritimum* showed increased lymphocytic infiltration and epidermal erosion, however bacterial cells were not present in the examined area (Figure 9B and 9C). The skin of fish exposed to *T. dicentrarchi* showed clusters of filamentous rods on the surface of the dermis (Figure 9D), and penetrating the dermis (Figure 9E) and underlying skeletal muscle (Figure 9F). One individual also showed extensive vacuolisation of the skeletal muscle (Figure 9D) along with increased lymphocytic infiltration.

**Figure 9.**
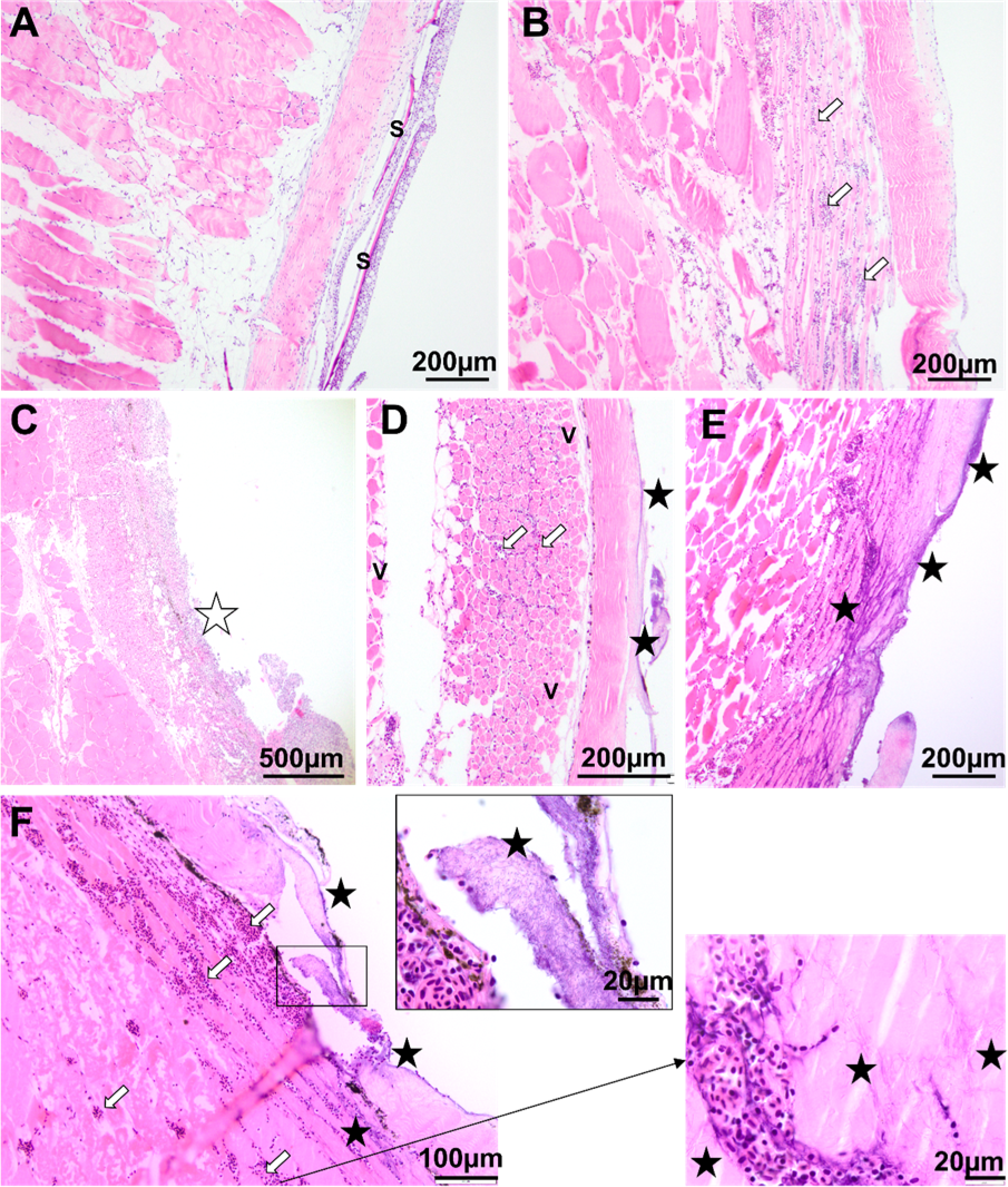
Histopathological changes in the skin of Chinook salmon, *Oncorhynchus tshawytscha*, exposed to *Tenacibaculum* species. A) Fish in the control group not exposed to *Tenacibaculum* spp. showing scales (S) and overlying epidermis. (B-C) Fish exposed to *T. maritimum*. (B) smolt exposed to *T. maritimum* showed increased lymphocytic infiltration (white arrows) and C) ulceration into the dermis (white star). (D, E & F) Fish exposed to *T. dicentrarchi*. (D) Vacuolization of skeletal muscle (V) and basophilic mats of filamentous bacteria (black stars) and lymphocytic infiltration (white arrows). (E and F) Filamentous bacterial mat (black stars) underlying skeletal muscle and increased lymphocytic infiltration (white arrows).

The gills of Chinook salmon showed a wide range of pathologies following exposure to *Tenacibaculum* spp. (Figure 10). Smolt exposed to *T. maritimum* and *T. dicentrarchi* displayed widespread, mild to severe lamellar hyperplasia, especially towards the base of the gills (Figure 10 C and 10D). There was local to widespread epithelial lifting of the secondary lamellae (Figure 10 D) and local to widespread and occasionally severe degeneration of the secondary lamellae (Figure 10 C, 10E and 10F). Thickening and adhesion of the secondary lamellae (lamellar fusion) were also observed, as were the presence of apoptotic erythrocytes within the secondary lamellae.

**Figure 10.**
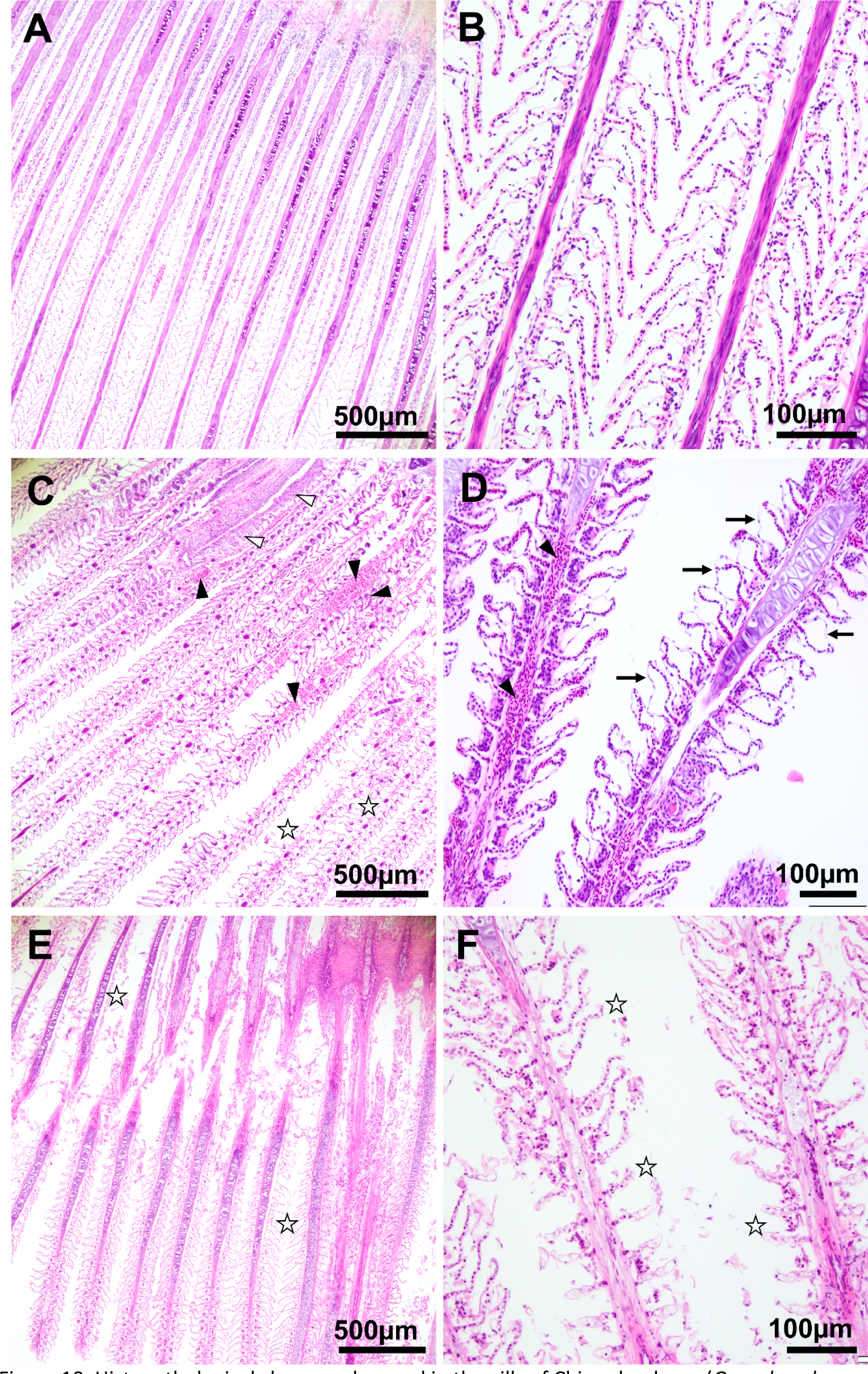
Histopathological changes observed in the gills of Chinook salmon (*Oncorhynchus tshawytscha)* exposed to *Tenacibaculum* spp. (A, B) Normal gills of Chinook salmon from the control group that was not exposed to *Tenacibaculum* spp. (C) Gill filaments of Chinook salmon exposed to *T. maritimum* showing primary lamellar hyperplasia (black triangles), complete degeneration of the secondary lamellae (white stars) and extensive lymphocytic infiltration (white triangles). (D) Gill showing further hyperplasia (black triangles) and epithelial lifting and necrosis (black arrow). (E, F) Extensive degeneration of the secondary lamellae (white stars) shown in salmon exposed to *T. dicentrarchi*.

### 3.4. Field evaluation of external vs systemic infection

Fish exhibiting gross signs of tenacibaculosis from marine farms showed *T. maritimum* growth by culture media from the surface of skin, eroded mouth, gills, and fins in 68% (28 out of 41 sampled; Table 2) of the population sampled. No culturable *T. maritimum* was isolated from any of the anterior kidney samples. *Tenacibaculum maritimum* DNA was detected in 9.7% (4 out of 41; Table 2) of infected fish although the kidney tissue appeared to be grossly normal with no culturable *T. maritimum*. This observation is consistent with the *in-vivo* challenge of *T. maritimum* (n = 29) and *T. dicentrarchi* (n = 30) in this study (Supplementary Table 1). Despite the low level of *T. maritimum* and *T. dicentrarchi* detection in the anterior kidney of challenged fish by ddPCR (Figure 7), no viable cultures were recovered from any of the corresponding anterior kidney samples of the challenged fish.

**Table 2.**
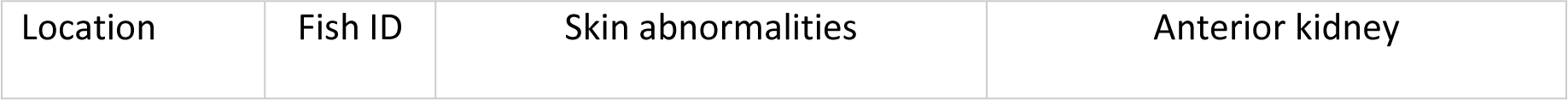
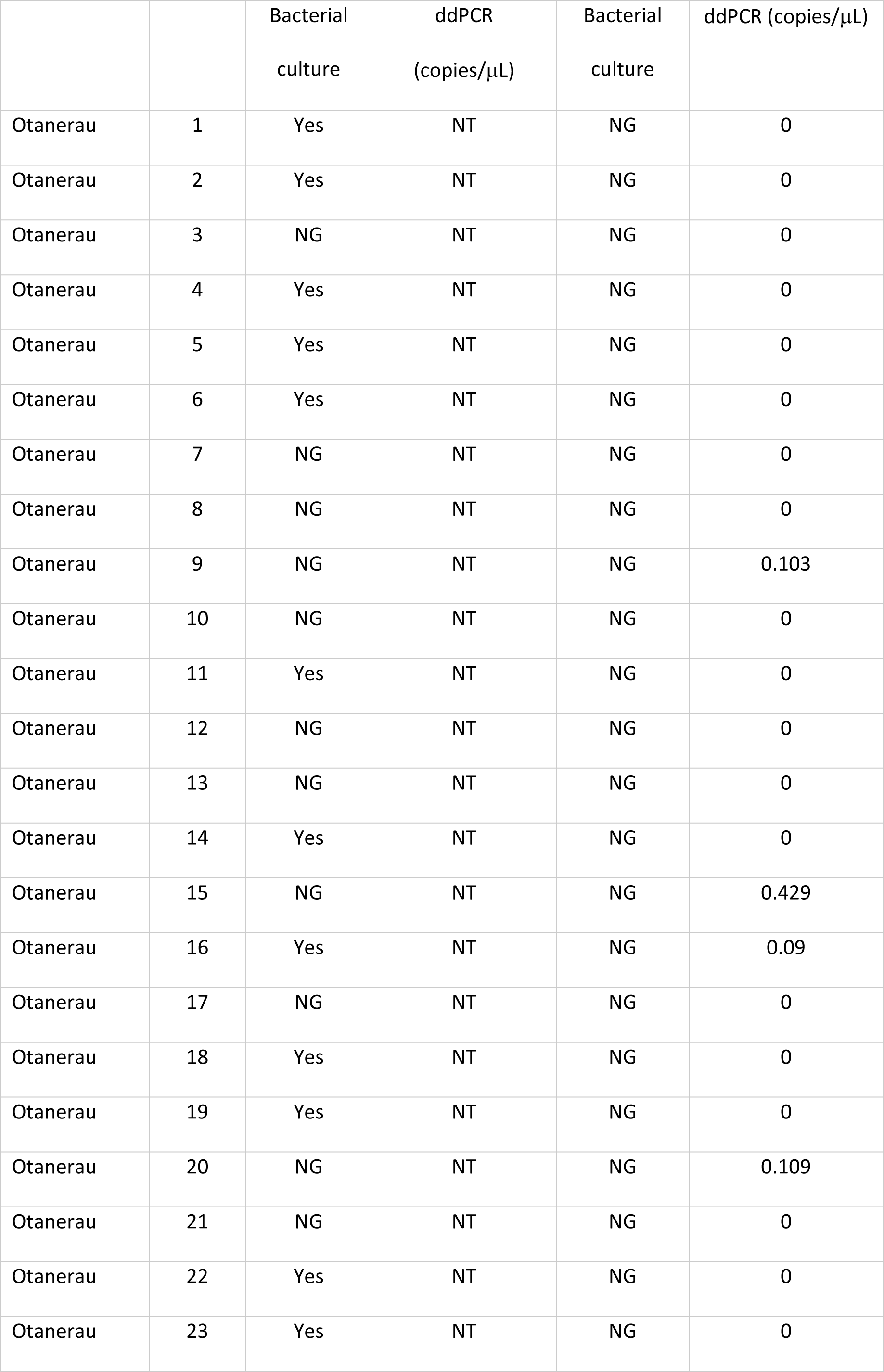
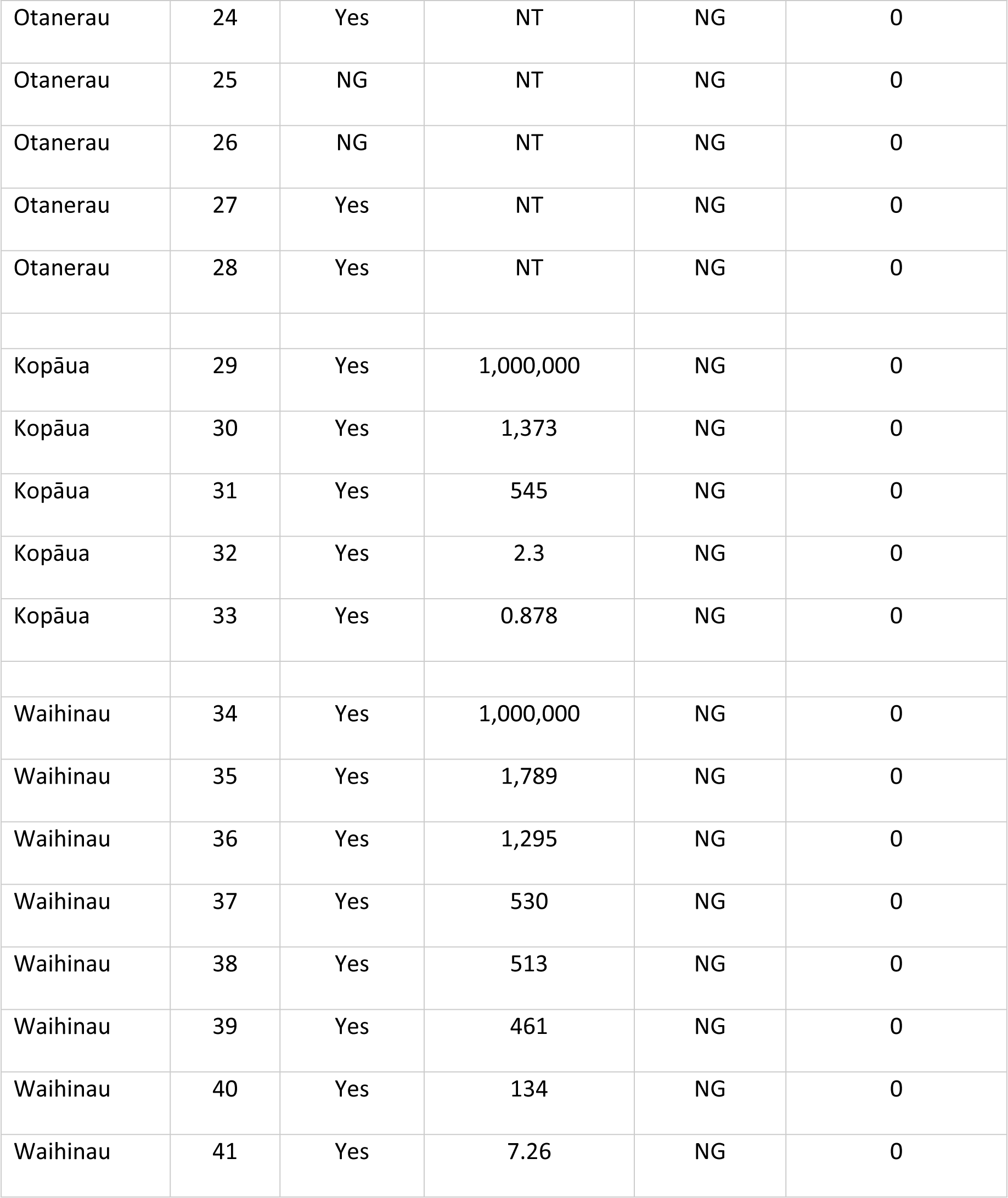
Viable and non-viable *Tenacibaculum maritimum* detection in skin and anterior kidney by bacterial culture and ddPCR in naturally infected Chinook salmon, all with skin lesions, from farm outbreaks. Yes = Growth of *T. maritimum*; NG = No growth of *T. maritimum*; NT = Not tested.

## 4. Discussion

Chinook salmon (*Oncorhynchus tshawytscha*) in Aotearoa New Zealand are susceptible to tenacibaculosis caused by the pathogens *Tenacibaculum maritimum* and *T. dicentrarchi*. In Aotearoa New Zealand, outbreaks of disease in Chinook salmon associated with skin infections were first reported in 2012 in sea pens located in the Marlborough Sounds and have been consistently reported thereafter [28, 31, 54, 55]. Assigning disease causation and confirming whether a ubiquitous microorganism is an infectious agent or is commensal flora of the host remains a challenge for diagnostic microbiologists, specifically for topical diseases [38, 56, 57]. Although most salmonids are susceptible to tenacibaculosis [15], it was reported that Pacific salmon species in the Pacific Northwest were resistant to mouth-rot disease caused by *T. maritimum* [see 8]. The absence of prior knowledge on the host-pathogen relationship between Chinook salmon and *Tenacibaculum* species, combined with the commensal bacterial populations [58], and NZ *Rickettsia*-like organisms [59] added to the complex diagnosis of tenacibaculosis in New Zealand Chinook salmon aquaculture. This is the first study to provide unequivocal evidence demonstrating the host-pathogen interaction between Chinook salmon and *Tenacibaculum* species. This study fulfils Koch’s postulates for the three molecular O-AGC types of *T. maritimum* (O-AGC Type 3-0, Type 2-1 and Type 3-2) and *T. dicentrarchi,* all of Aotearoa New Zealand origin, as pathogens of Chinook salmon by following a chronological approach: pathogen association and isolation, the cultured pathogen causes disease when introduced into a healthy organism [37, 60] and the counterfactual [60] that establishes causal evidence of *Tenacibaculum* species as a pathogen of tenacibaculosis in Chinook salmon. We confirmed that all molecular O-AGC types of *T. maritimum* and *T. dicentrarchi* isolated from Chinook salmon disease outbreaks in the Marlborough Sounds were shown to be highly infectious and induced skin disease in naïve, healthy fish in this study and that Chinook salmon farmed in Aotearoa New Zealand are susceptible to tenacibaculosis.

One of the difficulties in fulfilling Koch’s postulates is establishing a functional *in vivo* challenge model in a controlled laboratory setting which is essential to mimic the actual route of infection in the natural environment for a given pathogen [61]. Several experimental tenacibaculosis infection methods such as bath, intraperitoneal injection, prolonged exposure up to 18 h and injury induced infection have resulted in varying levels of success, including failure to reproduce tenacibaculosis [46, 61-65]. As Avendaño-Herrera et al. [24] described, the essential requirement of seawater for the growth of *T. maritimum* plays a critical role in achieving a successful culture and an *in vivo* challenge model. As shown in Figure 2, we obtained a homogenous growth of *T. maritimum* when it was cultured in MSM (with natural seawater) in comparison to ZoBell’s general purpose Marine Agar broth. We found that limiting the bacterial aggregation in broth is critical for achieving an accurate cell count and reproducible *in-vivo* challenge.

Although many marine bacteria have been successfully manipulated using artificial seawater [66-69], the intrinsic requirements for growth, viability and replication of *Tenacibaculum* species in natural versus artificial seawater is unknown. It is noted however, that an *in vivo* trial conducted using artificial sea water to test the efficacy of an autogenous bivalent vaccine against *T. maritimum* failed to reproduce tenacibaculosis [65], which suggests that natural seawater is a pre-requisite for achieving infection for *in vivo* experimentation. Furthermore, it is evident that failing to simulate the essential characteristics of the natural environment, may likely affect the microbial growth kinetics and physiology of the putative pathogen [70-74]. Immersion replicates the natural route of a non-vector-borne disease, whereby the pathogen must penetrate the host’s first line of defence, the mucus barrier [75, 76].

We conducted the experimental infection using a 100% natural seawater bath immersion as our challenge model and were able to achieve a successful induction of tenacibaculosis in naïve Chinook salmon. Although fish were exposed to pathogens for a short time (1 h), this was sufficient for the pathogen concentration used in this trial to establish infection with subsequent progression of the disease after the infected fish were transferred to a continuous RAS. Although a low level of *Tenacibaculum* spp. had been introduced to the RAS tanks during fish transfers from the immersion challenge bath, the daily reduction in viable cell count of *Tenacibaculum* spp. in RAS tank water suggests that the continuous UV irradiation was reducing the background load of *Tenacibaculum* spp. either due to replication or to shedding from infected fish. Progression of tenacibaculosis despite the declining *Tenacibaculum* load in tank water confirms the ability of these pathogens to adhere to fish skin, mucus or gills and to proliferate to cause disease after a short period of exposure [15, 77].

The pathologies observed in our experimental infection of *T. maritimum* and *T. dicentrarchi* are consistent with most clinical symptoms seen in Chinook salmon summer mortality in Aotearoa New Zealand [29, 31, 78] and tenacibaculosis observed in other salmonids [5-9, 16, 17, 79-82]. A compelling difference in the degree of skin and cartilage necrosis was seen between *T. maritimum* and *T. dicentrarchi* infection in this study. Fish infected with *T. maritimum* presented spreading but shallow skin necrosis whereas *T. dicentrarchi* infection caused deeper ulcers extending into the musculature, along with intensive cranial and caudal peduncle cartilage necrosis. Yellow plaques of bacterial mats on the fish abdomen and gills were commonly observed with both species of pathogen as described in other studies [8, 16, 62]. The re-isolation of the respective species from patches of bacterial mat and epidermal abnormalities (i.e., skin ulcer, fin necrosis and mouth rot) indicates the proliferation of the pathogens, their invasion of the mucus-epidermal layer and their ability to evade mucosal immunity [15, 77, 83, 84].

We noted a relationship between fish size and mortality, whereby there were survivors amongst the large fish challenged with *T. maritimum* O-AGC Type 3-0, but there were no survivors amongst the smaller fish challenged with *T. maritimum* strain O-AGC Type 2-1 and Type 3-2. These apparent differences in susceptibility would appear to be related to fish size rather than serotype. The mean weight of fish challenged with O-AGC Type 3-0 was 223 ± 4.5 g while for the other two serotypes the mean weight was 120 ± 1.2 g, which suggests either that smaller fish are more susceptible to tenacibaculosis or that there is inter-strain variation in virulence. Despite the different survival rates, all three molecular O-AGC types of *T. maritimum* were able to cause disease in Chinook salmon.

Bacterial colonisation was evident grossly in the skin and gills of both *T. maritimum* and *T. dicentrarchi* infected fish; however, only the skin of fish infected with *T. dicentrarchi* showed the presence of bacterial cells in histopathological analysis. Histologically, fish infected with *T. dicentrarchi* and *T. maritimum* exhibited a range of gill pathologies including lamellar hyperplasia and fusion which has been observed in Atlantic salmon with amoebic gill disease co-infected with *T. dicentrarchi* [see 85]. Despite the histologically evident gill pathologies in fish infected with *T. maritimum* and *T. dicentrarchi*, the absence of bacterial cells in histology sections of gills and *T. maritimum* infected skin could be due to the fixation process. Alternatively it suggests the potential involvement of bacterial toxic metabolites such as extracellular products (ECPs) in hydrolysing epithelial cells as seen in other disease caused by fish pathogens such as *Aeromonas hydrophila*, *Aeromonas salmonicida*, *Flavobacterium psychrophilum Photobacterium damselae* ssp. *damselae*, *Ph. Damselae* ssp. *piscicida*, *Streptococcus iniae,* and *Yersinia ruckeri* [see 86, 87-94]. As noted by van Gelderen et al. [95] the ECPs of *T. maritimum* were lethal and caused gill necrosis in experimentally challenged Atlantic salmon.

In a recent study, Escribano et al. [96] also described the virulence components of *T. maritimum* ECPs including proteolytic and hydrolytic enzymes such as chondroitinase, sialidase, sphingomyelinase, ceramidase and collagenase, which are associated with host tissue degradation. The same study also demonstrated the toxicity of *T. maritimum* ECPs on artificially infected *epithelioma papulosum cyprini* cell line and Senegalese sole (*Solea senegalensis)* fingerlings whereby the cytotoxic effect on the cell line and ulcerative haemorrhagic lesions on fingerlings were observed within 24 h post-challenge. The involvement of ECPs of *Tenacibaculum* species in causing tenacibaculosis requires further investigation because ECP based vaccination has shown protection against some important bacterial diseases including yersiniosis and streptococcosis [97-100].

Bacterial pathogenesis plays a key role in determining the mechanism and development of an infection [101] and this understanding is crucial for the development of a viable disease prevention option (e.g. vaccines) [15, 102]. There are a few studies reporting that *Tenacibaculum* spp. (primarily *T. maritimum*) cause systemic infection and have been detected in internal organs of Atlantic salmon and turbot (*Psetta maxima*) [see 8, 46]. However, tenacibaculosis is a topical skin disease that causes extensive damage to fish skin and cartilage [15] and its ability to cause septicaemia or internal infection is unclear if detection is solely based on presence of nucleic acids and where culture has not been used [23].

In Chinook salmon, we found *Tenacibaculum* species infect the skin, fins, and cartilage tissue; no substantial gross pathology was observed in the internal organs such as spleen, swim bladder, liver, and kidney (one exception where a *T. maritimum* challenged survivor exhibited necrotising kidney epithelial membrane). In our study, viable *Tenacibaculum* species were not re-isolated from the anterior kidney of farmed or experimentally challenged Chinook salmon. Given the average seawater osmolality is 1000 mOsm kg^-1^ [47, 103] while fish body fluid is 280-360 mOsm kg^-1^ [104], the likelihood of *T. maritimum* to infect and proliferate in fish internal organs is low due to its obligate requirement for seawater [105]. The significantly low level of *Tenacibaculum* spp. detected in the anterior kidney of farmed and experimentally challenged fish using ddPCR (Table 2 and Figure 7) suggests, although there were no viable bacterial cells in fish internal organs but non-viable cells or nucleic acid of this pathogens had gained entry to internal organ via haematogenous spread from cite of infection from the skin [106, 107]. The positive correlation noted in bacterial load between external tissue (skin and gill) and anterior kidney in *T. maritimum* challenged fish further proves the likelihood of hematogenous spread of pathogen. The anterior kidney in fish is a known lymphoid organ and a site for antigen presentation, therefore during a *Tenacibaculum* infection, phagocytic antigen presenting cells may also be responsible for the presence of nucleic acid in this tissue [108]. This appears consistent with the observations of Frisch, Småge (8), who found *T. maritimum* was detected by real-time RT PCR in several internal organs, however fish were dying with no gross internal abnormalities other than “yellow plaques” on the jaw in experimentally induced mouth rot in Atlantic salmon smolts. Based on these findings, we conclude that tenacibaculosis is an external disease, therefore *T. maritimum* and *T. dicentrarchi* do not cause septicaemia. However, the secretome produced by the viable bacteria present on the external infection site may cause a systemic effect [95, 96].

In conclusion, this study has fulfilled Koch’s postulates for the three molecular O-AGC types of *T. maritimum,* and for *T. dicentrarchi,* which are independently causative agents of tenacibaculosis in Chinook salmon. Disease was induced at temperatures typically experienced during Aotearoa New Zealand summers and without any physical intervention. The cumulative survival of 40 ± 20% and 72 ± 9.2% in *T. maritimum* (O-AGC Type 3-0) and *T. dicentrarchi* challenges respectively, showed that under our controlled test conditions, some fish naturally survived the bacterial infection, which may indicate some innate resistance. Indeed, whole genome quantitative trait locus (QTL) mapping of a few fish (families or species) have indicated the presence of resistance genes towards certain bacterial pathogens such as *Flavobacterium psychrophilum* in rainbow trout [109-112] and *Flavobacterium columnaris* in catfish [113]. Our natural disease challenge model could be used to explore disease resilient breeding for more sustainable production amid increasing seawater temperatures linked to directional climate change [114] and would also have significant value in assessing the efficacy of prototype *Tenacibaculum* vaccines.

## Acknowledgements

This project was funded by New Zealand Ministry of Business, Innovation and Employment Endeavour Programmes Aquaculture Health to Maximise Productivity and Security (CAWX1707) and Emerging Aquatic Diseases: a novel diagnostic pipeline and management framework (CAWX2207). Kumanan was supported by an Australian Government Research Training Program scholarship. We thank the following Cawthron colleagues for assistance with this study: Gareth Nicolson, Michael Scott, Daniel Cross, Juliet Butler, and Hannah Appleton for assistance with animal rearing, acclimation and recirculation system management; Ian Saldanha for assistance with animal ethics and animal welfare; and Mark Englefield for coordination and logistics for microbiological manipulation. Zac Waddington and Stuart Barnes (New Zealand King Salmon Co. Ltd) provided salmon smolts for the in vivo experiment. Eden Cartwright (Bird Circus) designed the graphics Figures, 1, 5 and 6. None of the authors have conflicting industrial links or affiliation.

